# Functional Characterisation of Recombinant Proteins Using Ion Channel Switch Technology: A Label-Free, Wash-Free Platform for Biotechnological Applications

**DOI:** 10.64898/2026.02.13.705684

**Authors:** Mohammad Pourhassan Moghaddam, Krishanthi Jayasundera, Lele Jiang, Bruce A. Cornell, Stella M. Valenzuela

**Author notes:** **Corresponding Author:** Dr Mohammad Pourhassan Moghaddam. 1.Graduate School of Biomedical Engineering, Faculty of Engineering, University of New South Wales, Sydney, NSW, Australia., 2. BioPoint Pty Ltd, Belrose, NSW, Australia.

## Abstract

We present a rapid, label-free, and highly sensitive platform for characterising recombinant protein functionality using Ion Channel Switch (ICS) technology. This method enables precise evaluation of binding specificity, oligomeric state discrimination, and real-time analyte detection, addressing key challenges in protein engineering and bioprocess quality control. Using engineered single-chain variable fragments (scFv), ICS reliably distinguishes monomeric from multimeric forms, facilitates wash-free detection of analytes ranging from small molecules to larger biomolecules, and enables quantitative biosensing within seconds in real-time and continuous format. These capabilities establish ICS as a powerful tool for streamlining recombinant protein screening, with broad applications in diagnostics, therapeutic quality control, and automated bioprocess workflows.

## 1. Introduction

Recombinant proteins play an essential role in biotechnology, with applications spanning diagnostics, therapeutics, vaccines (Tripathi and Shrivastava, 2019), and industrial processes (Puetz and Wurm, 2019). Accurate and efficient functional characterisation of recombinant proteins is paramount for assessing their performance in these applications, ensuring structural integrity, and validating specific interactions. Traditional techniques for characterising recombinant proteins, such as Enzyme-Linked Immunosorbent Assay (ELISA), Surface Plasmon Resonance (SPR), and Size-Exclusion Chromatography (SEC), often rely on time-consuming protocols, require labelling agents, or are limited in their ability to distinguish subtle conformational differences between protein states. ELISA provides good sensitivity but may not capture the real-time interactions between the antibody and the antigens of interest (Baker et al., 2002). Surface plasmon resonance (SPR) enables real-time analysis of molecular interactions; however, it can be affected by nonspecific binding and demands meticulous surface preparation (Stahelin, 2013). Size-exclusion chromatography (SEC) effectively separates proteins based on size (Baker et al., 2002), though it may not detect minor conformational differences, nor does it provide information regarding molecular interactions.

Ion Channel Switch (ICS) technology presents a transformative solution to these challenges. ICS is a biosensor technology that measures real-time conductance changes in tethered bilayer lipid membranes (tBLMs) containing gramicidin A (gA) ion channels. This system enables sensitive, rapid detection of analyte binding events even in complex sample matrices. This platform has the potential to offer the advantages of methods such as ELISA, SPR, with the capability of performing real-time interaction measurements (Cornell et al., 1997). The current study investigates the utility of ICS for screening recombinant single-chain variable fragments (scFvs), highlighting its capabilities in evaluating protein functionality, specificity, and quantifying analyte interactions in real time. Furthermore, the study demonstrates the application of recombinant proteins in developing a continuous label-free biosensing method.

## 2. Materials and Methods

### 2.1 Recombinant Protein Design, Purification and Expression using Mammalian Cell System

A scFv, derived from an anti-P4 antibody (Dubreuil et al., 2005), was adopted as the main P4-binder component, and an irrelevant scFv, derived from an anti-human ferritin antibody was chosen as the non-binder protein control (Nymalm et al., 2002; “US20090074657A1 - Nucleotide and protein sequences of an antibody directed against an epitope common to human acidic and basic ferritins, monoclonal antibodies or antibody-like molecules comprising these sequences and uses thereof - Google Patents,” n.d.). A streptavidin (SA) domain (Kroetsch et al., 2018) sequence was fused upstream of the scFvs sequence. SA is a well-known protein that exhibits a high affinity for biotin (Sano et al., 1998). This fusion was intended to create a binding site for the biotinylated elements on the biosensor surface. An amino acid linker was used to connect the SA domain to the scFv sequences (to produce SA-antibody proteins), ensuring proper spacing and flexibility between these functional domains. Final sequences are presented in Table 1.

**Table 1.**
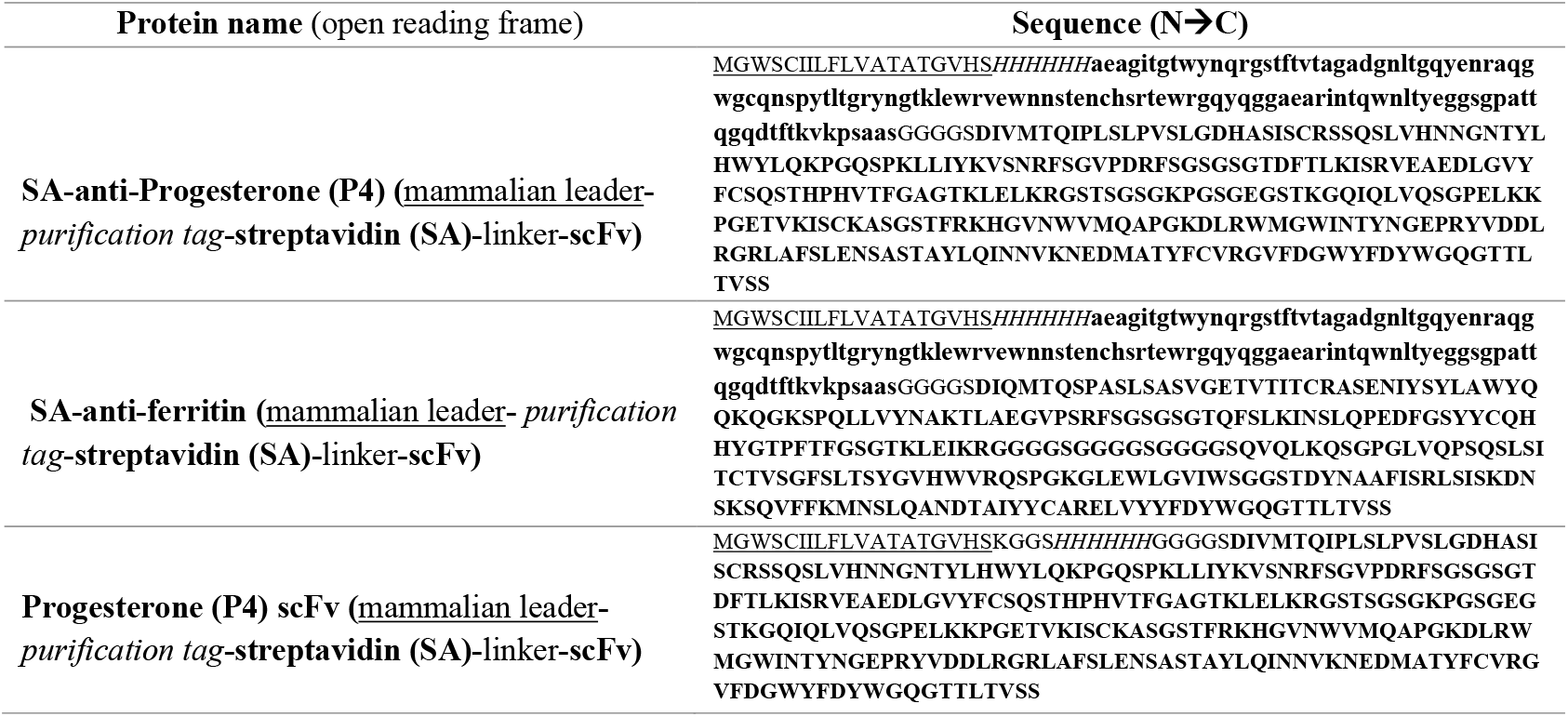
The final designed sequences of the analyte binders.

Recombinant constructs encoding SA–antibody fusions and the scFv were codon-optimised for expression in CHO cells and synthesised by Geneart (Thermo Fisher Scientific). Each gene was cloned into the pcDNA3.1 vector, transformed into *E. coli* Sigma Sig10 competent cells (Sigma-Aldrich), and cultured under standard conditions. Plasmid DNA was purified using the NucleoBond Xtra Midi Kit (Macherey-Nagel, Cat# 740410.100), eluted in ultrapure distilled water (Invitrogen, #10977-015), and washed with absolute ethanol (Merck, #1.00983.2511). All steps were carried out under sterile conditions.

ExpiCHO™ cells (Thermo Fisher Scientific) were maintained in ExpiCHO™ Expression Medium (#A29100-01) at 37 °C with agitation at 130 rpm, 80% humidity, and 7.5–8.0% CO_2_. Transient transfections were performed using the ExpiFectamine™ CHO Transfection Kit (#A29129) according to the manufacturer’s High Titer protocol. Cultures for SA– antibody fusions were scaled to a total volume of 1 L in Erlenmeyer flasks (Corning, #431145), while a 100 mL culture was used for scFv expression. At 20–22 hours post-transfection, cultures were supplemented with ExpiCHO™ Feed (#A29101-02) and Enhancer (#100033018), followed by incubation at 32 °C. On Day 6, cells were harvested by centrifugation. Supernatants were clarified using 0.2–0.22 µm PES filters (Corning; Thermo Scientific Nalgene #568-0020, #564-0020) or syringe filters (Millipore) and stored at −80 °C until purification.

Purification was conducted using an A KTA Pure system (GE Healthcare) and immobilised metal affinity chromatography (IMAC). Five millilitre columns (HiTrap Excel, Cat# 17-3712-05; HisTrap Excel, Cytiva) were equilibrated with DPBS (Lonza) for SA-fusions and 20 mM sodium phosphate (Na_2_HPO_4_) with 500 mM sodium chloride (NaCl) for scFv purification. Clarified supernatants (approximately 920 mL for SA-fusions and 100 mL for the scFv) were loaded onto the columns, and bound proteins were eluted with 500 mM imidazole. Eluted proteins were buffer exchanged into DPBS using HiPrep 26/10 desalting columns (GE Healthcare, Cat# 17-5087-01) or 30 kDa MWCO Amicon Ultra-15 centrifugal filter units (Millipore).

Final protein concentrations were adjusted to 1–2 mg/mL, sterile filtered through 0.22 µm membranes, and preserved with 0.05% sodium azide (10% stock in PBS). Proteins were formulated in 50% glycerol (100% stock) for storage. Concentrations were determined using a NanoDrop 2000 UV spectrophotometer (Thermo Scientific).

### 2.2. SDS-PAGE Analysis of Purified Recombinant Proteins

The molecular size of the samples was analysed using SDS-PAGE, followed by Coomassie blue staining (Gallagher (electrophoresis, staining) and Sasse (staining), 1998). Briefly, 100ng/ul of the protein samples were mixed with Laemmli buffer in a 1:1 ratio, heated for 15 minutes at 95◦C, and 30ul of the heated samples were loaded into pre-cast gels (MINI-PROTEAN, 4-15% Catalog # 4568084, Bio-Rad), including a protein size marker (Catalog # 161-0373, Bio-Rad). The gel was run for 40 minutes at 140 V in 1x reducing SDS-PAGE using Tris/Glycine/SDS buffer (Catalog #1610771, Bio-Rad). The gel cassettes were carefully opened, and the exposed gels were rinsed with Milli-Q water, followed by staining with Coomassie blue solution (0.2% Coomassie G250 dye in the solution of 10% acetic acid and 40% methanol) for 30 minutes, and destained with the destaining buffer (10% acetic acid and 40% methanol) and paper absorbents for 3 hours while shaking. The destained gels were rinsed with milli Q water. The gels were photographed using gel doc (ChemiDoc MP imaging system, Bio-Rad).

### 2.3. Functional characterisation of the expressed proteins using Surface Plasmon Resonance (SPR)

In order to test the functionality of the expressed proteins, Surface Plasmon Resonance (SPR) (Biacore T-200, Cytivalifesciences) was used to characterise the binding of the proteins to P4 and or biotin. For this reason, BSA-P4 (Catalog # 80-IP50, Fitzgerald), BSA-biotin (Catalog # A8549-10MG, Sigma Aldrich), or BSA (Catalog # 23209, Thermo Scientific) were conjugated onto individual CM-5 chips (Cytivalifesciences). All experiments were performed using degassed and pre-warmed buffers as recommended. CM-5 chips, which carry a matrix of carboxymethylated dextran covalently attached to a gold surface and have four channels, were used to immobilise BSA-P4, BSA-biotin, and BSA (as negative control) proteins in separate channels of the chip via EDC/NHS chemistry in-situ, following the recommended procedure from Cytiva Life Sciences. Sodium Acetate (10 mM, pH 5) was used as conjugation buffer, and the concentration of the BSA proteins was kept at 20 μg/ml for each conjugation. The first channel was kept unmodified to serve as the reference channel. The expressed protein samples (non-blocked and biotin/P4 blocked samples) were diluted in the HBS-EP+ running buffer (0.1 M HEPES, 1.5 M NaCl, 0.03 M EDTA and 0.5% v/v Surfactant P20) and injected into the system and the binding events were monitored. As another control, SA-antibody proteins were incubated with their corresponding ligands and used as non-binding controls.

### 2.4. ICS Sensor Preparation and Testing

tBLM sensors were fabricated following the literature10. The SDx gold-coated slides (SDx Tethered Membranes Pty Ltd), possessing six sensing electrodes and pre-coated with tethered first layer lipid on top of the electrodes, were bonded with a flow cell top-on cartridge (SDx Tethered Membranes Pty Ltd), pre-coated with the gold counter electrode and a double-face adhesive laminate layer covering the electrodes: the gold slide was placed and aligned in a special alignment jig, leaving the top part facing up. Then, the top layer of adhesive was removed from the cartridge and was attached to the gold slides, ensuring the correct orientation of the slide with both the jig and the cartridge by following the marks on the jig, the slide and the cartridge. Then, a silicon rubber pressure pad and an aluminium pressure plate were aligned on top of the attached cartridge, and the assembly was inserted in a press clamp. Then, a minimum of two-minute pressure was applied to the assembly by closing the knob of the clamp to maximise the bonding between the slide and the cartridge. After releasing the knob, the assembly was taken out of the clamp and dissembled gently to have the bonded electrode-flow cell cartridge assembly ready for the next step. Immediately, the second lipid layer was created; 8ul of an 3mM ethanolic solution, containing 70 %:30 % molar mix of diether diphytanyl (C16) phosphatidyl choline:diether diphatanyl (C16) hydroxyl (SDx Tethered Membranes Pty Ltd) and gramicidin A-P4 or gramicidin-biotin, was added to each well, starting from well one with a 10 s incubation time between each addition. After the last well was loaded with the second layer solution and after one minute incubation, 150 μl PBS was added to each well, starting from well one with a 10 s incubation time between each addition. Then, all wells were rinsed simultaneously three times. The ready biosensor (figure 3.2) was plugged into the EIS reader for performing the experiments. An amplitude of 25 mV was used as the input potential with the scanning range of 2 Hz to 2000 Hz for the EIS step. Before the materials were added for experiments, the membrane conductance, i.e. Gm, and membrane capacitance, i.e. Cm, were monitored. As a quality control step, the assembled chips that had Cm around 20 nF were kept for the experiments because Cm∼20 nF indicates a successfully formed lipid bilayer. The EIS reader software, called SDx TethaQuick (SDx Tethered Membranes Pty Ltd), was used to capture the EIS data output.

After assembly of the ICS biosensor, anti-P4 and anti-ferritin proteins, as the negative control, were added to the sensor to study their functionality using the biosensor. For this reason, different membranes containing different gA conjugates were used: a tBLM containing only gAP4, a tBLM containing only gA-biotin and a tBLM containing gA-P4 and a lipid-biotin or a gA-biotin. After fabrication of the ICS cartridge containing the above tBLMs, 150μl of PBS was removed from the well, 50μl of the expressed protein was added to the wells, and any decrease in the membrane conductance was monitored using tetaQuick software.

For validation of the P4 ICS mechanism, response of the P4 ICS was studied against different controls and experimental conditions, including the use of: binding and non-binding proteins, different analytes, different gA modifications, a tBLM without gA, different pH and varied incubation times. BSA, BSA-P4, and P4 were used as the analytes to study the ICS response to addition of different analytes. To test the effect of changing gA modification, in addition to the gA-P4 and gA-biotin., gA modified with either hydroxyl (-OH) or tert-butyloxycarbonyl (-BOC) chemical groups were employed as the gA controls. The effect of changing pH on the P4 ICS function was also investigated; for performing the pH experiments, after assembly of the cartridge and connection to the EIS reader, the tBLM was rinsed at least three times with a buffer of specific pH. Then, the anti-P4 protein in the same buffer was added to the tBLM. The P4 ICS response in the pHs of ∼4.7, 5.8, and 10 was investigated.

To distinguish monomeric and multimeric forms of scFvs, a sequential binding strategy was employed. Pre-incubation with a monomeric binder blocked secondary binding by a multimeric competitor, allowing functional discrimination via signal suppression.

### 2.5. Detection of analytes with ICS biosensor

For testing the capability of the proposed P4 ICS for quantitative measurements, the biosensor was challenged using various concentrations of P4, BSA-P4 and SA-anti-P4. Briefly, after the fabrication of biosensors, 50μl of analytes in various concentrations were added to wells. After that, the biosensor signal reached a steady state, followed by rinse with 150μl PBS four times. For the competitive detections, various concentrations of P4 and BSA-P4 were added to the biosensor wells containing pre-added SA-anti-P4 of fixed concentration (120 nM), and the changes in the membrane conductance were registered for each case. A wash-free assay was developed using simultaneous application of recombinant binder and analyte. Conductance changes were measured to reflect competitive displacement, enabling rapid and accurate analyte quantification.

### 2.6. Quantitative analysis of biosensor signals

Membrane conductance signals were analysed by assessing the signal rate (K), signal size (span), and signal slope using GraphPad Prism version 9 (La Jolla, California, USA). The signal size (span) refers to the difference between the primary and secondary levels of membrane conductance after analyte binding. The signal rate describes the speed at which membrane conductance changes following analyte binding. Equilibrium represents the maximum signal size that stabilizes after analyte addition, while pre-equilibrium refers to the phase before the signal reaches its maximum.

To calculate the signal rate and span, normalized membrane conductance data were fitted using non-linear regression. For the SA-anti-P4 data, a one-phase decay model was applied, while for the P4 data, a one-phase association model was used to obtain equilibrium signal size and signal rate. Additionally, the slope of the pre-equilibrium signals was calculated using the first two data points after analyte addition. This allowed the determination of the minimum time needed to observe differences in signal levels at varying analyte concentrations.

The pre-equilibrium-based method provides a quicker response, typically within one minute. ICS signals were quantified using both equilibrium and pre-equilibrium methods. Linear response curves for P4, BSA-P4, and SA-anti-P4 were generated to assess concentration-dependent responses.

### 2.7. Molecular modelling and docking simulations

The tertiary structure (3D) of SA-anti-P4 was predicted using DeepMinds Alphfold (Jumper et al., 2021), which is considered one of the most accurate tools for protein structure prediction. The structural predictions were implemented via ColabFold python-based online server using the AlphaFold MMseqs2 algorithm, including an AMBER relaxation step to relax the predicted structures (Mirdita et al., 2022).

The structure of P4-C3 linker was drawn in the ChemDraw version 20 (PerkinElmer Informatics) and was converted to the SDF format, and the SDF formats of P4 and P4-C11 linker were downloaded from the PubChem database. CB-Dock tool was adopted to dock SDF files of P4 and the P4 linker molecules to the predicted 3D structure of SA-anti-P4 following the default parameters. After docking, the ligand poses with the lowest docking score was selected as the binding pose for the ligands to the designed proteins (Liu et al., 2020). The complexes of docked ligands-SA-anti-P4 were visualised using PyMOL molecular visualisation tool (Schrodinger LLC, 2015), where a reference P4-anti-P4 Fab complex (PDB code: 1dbb) was used to confirm the docked poses of P4 and P4-linker structures on SA-anti-P4 protein.

### 2.8. Continuous Biosensing

Given that the SA-anti-P4 protein possesses streptavidin and anti-P4 scFv domains, the possibility of repeated P4 detection was tested using tBLMs containing only gAP4 (pure gAP4) and a mixture of gAP4 and gAbiotin. Briefly, the second layer containing either 50 nM gAP4 or mixture of gAP4 and gAbiotin was prepared. 120 nM SA-anti-P4 protein was added to each tBLMs type, and the gating signal was followed. After reaching the signal to plateau, the tBLMs were rinsed 4x with PBS (the baseline signal), followed by addition of 10 nM P4. The signals were again monitored to reach the plateau, and the wells were rinsed with PBS. When the signal declined to the baseline level, the next rounds of P4 addition and rinse were conducted.

## 3. Results and Discussion

### 3.1 Recombinant Protein Expression and Characterisation

SDS-PAGE analysis confirmed that the recombinant proteins were expressed at the expected molecular weights (Fig. 1A).

**Fig. 1.**
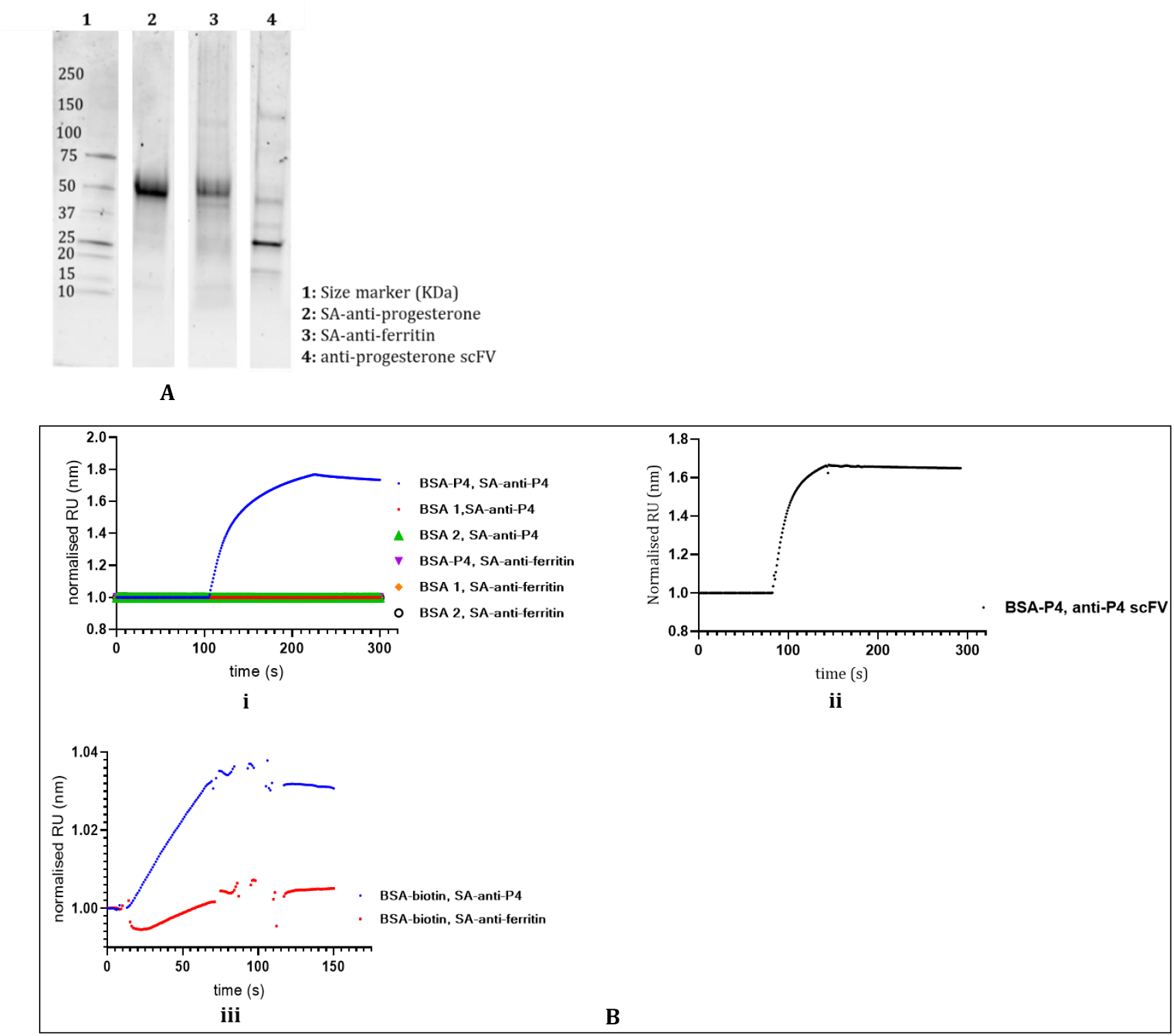
Characterisation of the expressed proteins. (A): SDS-PAGE analysis of the expressed recombinant proteins. The molecular weight of the proteins was found to be similar to the computationally predicted values provided by ProtParam tool (Table 2). (B): Functional analysis of the proteins using SPR. The SPR experiments confirmed the functionality and specificity of the expressed proteins. Anti-P4 proteins effectively bound to BSA-P4, while the anti-ferritin protein did not bind to BSA-P4. SA-anti-P4 showed binding to both BSA-P4 and BSA-biotin, but not to BSA alone (i and ii). Both SA-anti-P4 and SA-anti-ferritin bound to BSA-biotin, albeit with different binding levels (iii).

### 3.2 Binding Specificity

Below, SPR and ICS experiments demonstrated that the engineered scFvs specifically bound to their targets. Anti-P4 scFvs exhibited strong binding to P4-modified surfaces, while anti-ferritin scFvs showed no binding to these surfaces, confirming the specificity of the scFvs.

**Table 2.** Calculated physicochemical characteristics of the proteins.

| protein | Length (aa) | MW (KDa) | Isoelectric point |
| --- | --- | --- | --- |
| anti-progesterone scFv | 265 | ~29 | 9.2 |
| SA-anti-progesterone | 380 | ~41.6 | 8.89 |
| SA-anti-ferritin | 369 | ~39.8 | 8.61 |

#### 3.2.1. Functional characterisation of the expressed proteins using SPR

In the SPR experiment, the expressed proteins were used as test samples, while anti-ferritin, BSA, and SA-antibody proteins pre-blocked with P4 and/or biotin were employed as the negative control sets. SPR experiments demonstrated that both SA-anti-P4 and anti-P4 scFv proteins bind to the BSA-P4. However, neither the anti-ferritin nor the P4-blocked SA-anti-P4 proteins could bind to the BSA-P4. None of the tested proteins bound to BSA, indicating good specificity, where even the anti-P4 proteins, which had their parent antibody raised against the BSA-P4 antigen, were capable of binding to the P4 molecule only and not to BSA. In addition, the SA-antibody proteins were shown to bind to BSA-biotin. Blocking of the SA-antibody proteins with biotin abolished their binding to the BSA-biotin, confirming the functionality of the SA domain (Fig. 1B).

According to the SPR results, the designed bispecific SA-anti-P4 and the monospecific anti-P4 proteins are both capable of recognising P4, while the bispecific SA-antibody proteins can bind to biotin via their SA domain. P4 and biotin are also present in the surface layer of the ICS biosensor. Furthermore, blocking the P4 and biotin-binding sites by using free P4 or biotin cross-validated the functionality of SA-anti-P4 and SA-anti-ferritin. These experiments confirmed that the proteins were biologically active and suitable for testing and use in the ICS biosensor.

#### 3.2.2. Testing the specificity of analyte binding

The first step in analysing the recombinant proteins with the ICS biosensor was to test their binding specificity to gA channels embedded in the tBLM. gA channels modified with either hydroxyl (gA-OH) or tert-Butoxycarbonyl (gA-BOC) were tested as negative controls. The proteins were also added to tBLMs without gA channels. The results showed no biosensor signal, indicating no non-specific binding to gA or tBLM surface (Supplementary Figure 1). In the next step, we tested specific binding of the recombinant SA-antibody proteins to P4 or biotin conjugated on gA channels (i.e. gA-P4 or gA-biotin) (Fig. 2).

**Fig. 2.**
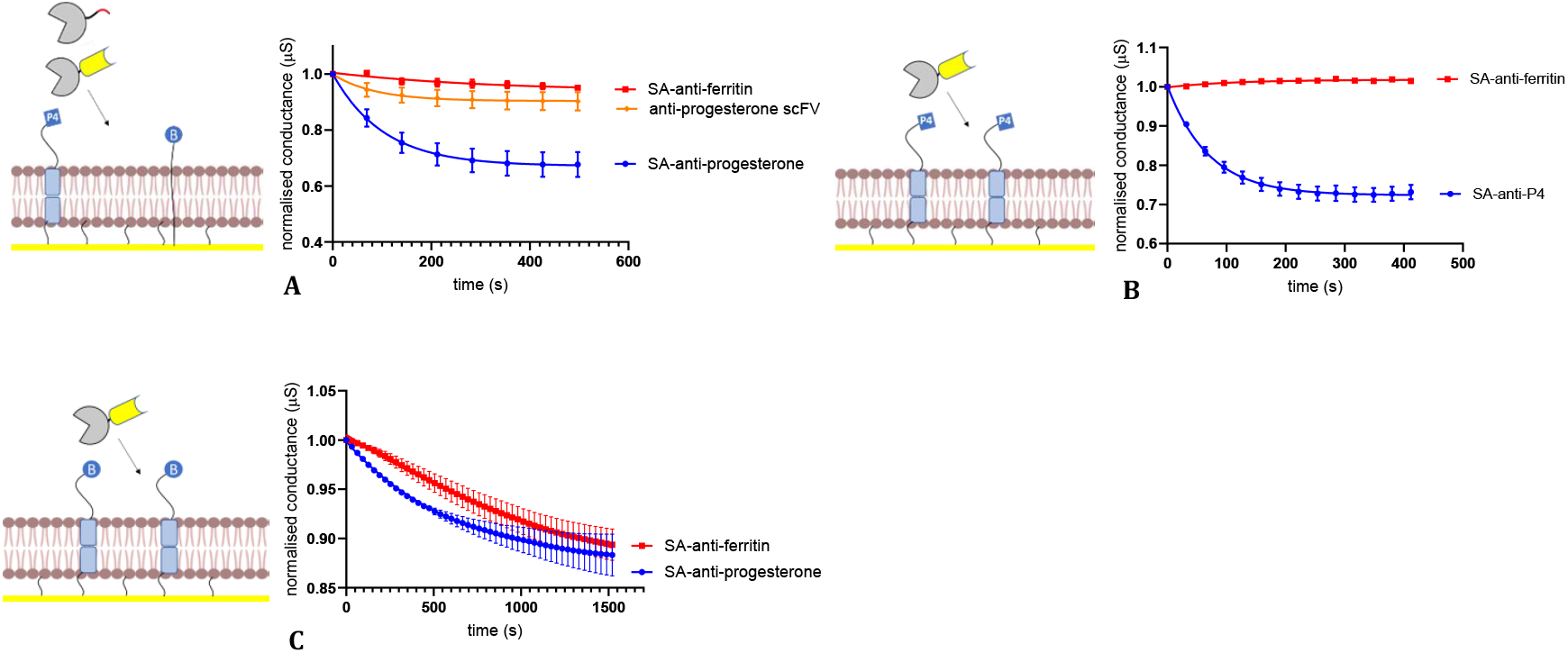
Characterisation of the expressed proteins using the P4 and biotin modified ICSs (n=3). (A): Testing the proteins using a tBLM containing a biotin presenting element such as a membrane-spanning lipid-biotin and the gA-P4 elements showed that SA-P4 protein could elicit a clear reduction in the membrane conductance. anti-P4 scFv and SA-anti-ferritin did not show a clear signal. Similarly, (B): for SA-anti-P4 a reduction in membrane conductance was observed in the membrane containing only gA-P4 too. (C): When the tBLM contains only gA-biotin, we observed a signal for SA-anti-progesterone and SA-anti-ferritin proteins.

Characterisation of the expressed proteins using P4 and biotin ICSs showed that in a tBLM containing membrane-spanning lipid–biotin and gA–P4, the SA–P4 protein induced a significant reduction in membrane conductance, whereas the anti-P4 scFv produced only a minor response, and the SA–anti-ferritin protein had little/no effect (one way ANOVA, P<0.0001) (Figs. 2A). A similar significant conductance reduction was observed in membranes containing only gA–P4 for SA-anti-P4 protein (paired t-test, P<0.0001) (Fig. 2B). As a negative control, SA–anti-ferritin did not gate gA–P4, confirming specificity. When the tBLM contained only gA–biotin, both SA–anti-progesterone and SA–anti-ferritin produced signals, indicating interaction with the biotin end of gA–biotin (Fig. 2C). A comparison between SA and scFv domains revealed differences in binding kinetics: the scFv domain from SA–anti-P4 exhibited a faster binding event than the SA domains in both SA–antibody proteins. Application-wise, this faster interaction kinetics may be advantageous in rapid biosensing, as the detection of target analytes would require less time.

Regarding the design of the SA–anti-P4 fusion protein, it was initially anticipated that no gating signal would be observed in membranes containing only gA, in the absence of biotin-presenting components. However, the detection of signals in membranes containing either gA–P4 or gA–biotin suggests that non-monomeric forms of the expressed fusion proteins may be contributing to the observed activity. To explore this possibility, we conducted a series of additional experiments and undertook a more detailed evaluation of the produced proteins.

### 3.3 Monomer versus Multimer Differentiation with ICS biosensing

Given the structure of the SA–anti-P4 protein as a bifunctional molecule, one would expect to see a decrease in membrane conductance only in the presence of biotin on the tBLM, through interaction between its streptavidin domain and the membrane-bound biotin. It was therefore not expected to induce gating in tBLMs containing only gA–P4 or gA–biotin. (Fig. 2). On the other hand, surface plasmon resonance (SPR) experiments (Section 3.2.1; Fig. 1B) confirmed similar P4-binding capability among all designed anti-P4 proteins. It is likely that the scFv bound to the P4 moiety of gA–P4 with comparable affinity to the SA– anti-P4 protein, but its interaction generated significantly smaller or no conductance changes. Therefore, to confirm that the observed signals were due to specific binding to gA–antigen molecules, a competitive inhibition assay was performed using the P4 ICS. Pre-incubation of anti-P4 scFv to a tBLM containing gA–P4, inhibited the response normally elicited by subsequent SA–anti-P4 addition, and the level of inhibition was dependent on the concentration of the pre-incubated scFv (Fig. 3A). In contrast, pre-incubation with SA–anti-ferritin had no effect on subsequent SA–anti-P4 activity, as shown by the expected decrease in conductance following its addition (Fig. 3B). These findings confirm that pre-binding of SA–anti-P4 blocked access to the gA–P4 sites for subsequent multivalent competitors. This outcome enabled discrimination between monomeric and multimeric scFv forms, highlighting the ICS system’s sensitivity to and ability to discern differences in protein structure. A proposed binding mechanism based on the inhibition studies is shown in Fig. 3C.

**Fig. 3.**
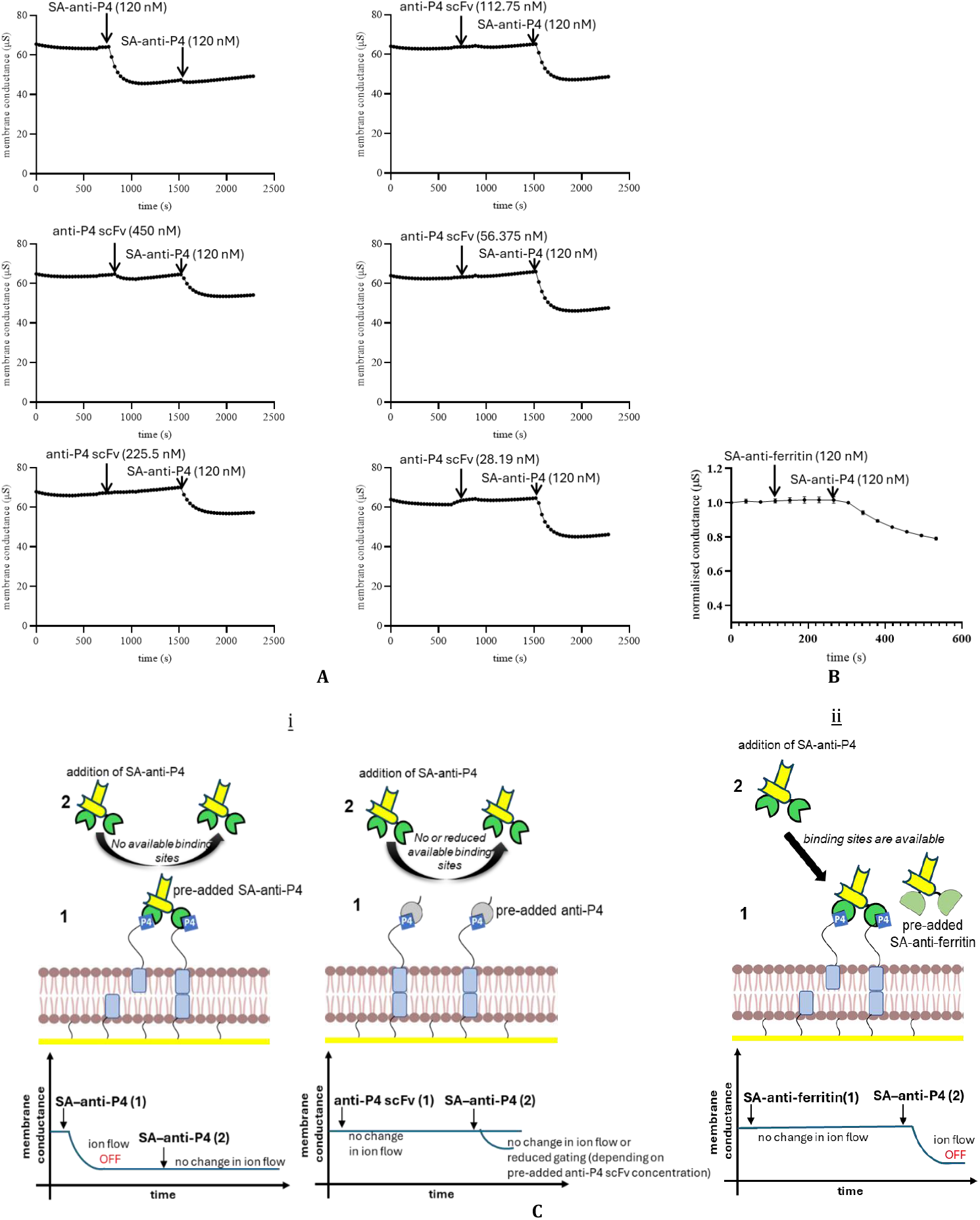
Monomer versus Multimer Differentiation with ICS biosensing. (A): Detection of a mostly monomeric binder by pre-incubating it with tBLM containing gA-analyte, such as gA-P4, followed by addition of the non-monomeric binder. If the monomeric binder binds to the antigen-modified gA, the secondary binder is not able to bind or have a reduced binding capapbility, as the specific binding sites on the modified gA are already occupied by the primary monomeric binders. (B): pre-addition of SA-anti-ferritin did not prevent SA-anti-P4 from the binding to gAP4. This is because the specific binding sites are available for binding to the SA-anti-P4 (n=3). (C): (i): Summary of the binding inhibition mechanism used to distinguish monomeric from non-monomeric gA–analyte binders. When gA–analyte sites are occupied by an initially added binder, no gating signal is observed upon adding a second binder. (ii): In contrast, if the first binder does not occupy the sites—due to low affinity or monomeric structure—a subsequent multivalent binder can still trigger a gating response. These results suggest that SA– antibody proteins are largely non-monomeric, while anti-P4 scFv is primarily monomeric. Aggregates inferred from ICS gating behaviour; not directly confirmed by native-state analysis.

The inhibition assay results confirmed that binding can occur without generating a significant gating signal, as observed with monomeric anti-P4 proteins. These findings indicate that binding to antigen-modified gA, such as gA–P4, and preventing re-association of gA monomers between the membrane leaflets is necessary for gating to occur. A monomeric binder with a single binding site can interact with gA–analyte molecules but cannot bridge multiple gA units diffusing in the upper membrane leaflet, and therefore does not induce a gating response. In contrast, SA–antibody proteins, likely possessing non-monomeric structures, produced significant gating signals. These results suggest that at least two binding sites per binder are required to elicit a gating signal in the ICS, which is consistent with previous findings where a biotinylated Fab (bFab) was used to cross-link antigen-modified gA molecules via streptavidin-bound MSL–biotin (Woodhouse et al., 1999). In this study, millimetre-scale electrodes with tethered bilayer lipid membranes (tBLMs) containing millions of ion channels were used. Conductance changes—arising from channel formation and dissociation—were monitored using electrochemical impedance spectroscopy (EIS). Multimeric antigen-specific binders, such as dimeric scFvs, reduce membrane conductance by pairing channels, unlike monomeric scFvs, which bind only one antigen-modified channel. This distinction enables the Ion Channel Switch (ICS) method to differentiate between monomeric and non-monomeric binding events.

One potential application of this technology is detection of non-monovalent immunoglobulins, such as IgE involved in progesterone (P4) hypersensitivity—a condition currently diagnosed using low specificity and low sensitivity skin tests (“Progestogen hypersensitivity - UpToDate,” n.d.). The system can also be used to study reversible binder associations; for instance, associated binders reduce conductance, while dissociation—potentially drug-induced—restores it. This suggests ICS may be useful for screening drugs that modulate protein-protein interactions (PPIs), a key therapeutic target (Petta et al., 2016; Thangudu et al., 2012). Additionally, ICS has potential for monitoring self-association in recombinant proteins, offering a faster and simpler alternative to techniques such as size exclusion chromatography and ultracentrifugation (Gell et al., 2012). Finally, ICS could be adapted to detect pathological protein associations, such as amyloid-β (Aβ) oligomers implicated in Alzheimer’s disease. Unlike ELISA, ICS has the potential to provide a rapid method for early Aβ oligomer detection (Wang et al., 2017).

### 3.4 Using the analyte binding recombinant protein in wash-free analyte detection

#### 3.4.1. Binding competition

The following section presents the competitive detection of P4 using the P4 ICS mechanism with the expressed SA–anti-P4 protein in a rapid, two-step, wash-free format. Detectable signals were observed within 15 minutes, demonstrating the efficiency of the assay. Both free and BSA-conjugated forms of P4 were tested. Specific responses were recorded for P4 (Fig. 4A) and BSA–P4 (Fig. 4B), while no signal was detected in response to unrelated analytes such as BSA. The control protein, SA–anti-ferritin, did not induce any response before or after the addition of P4 or BSA–P4, confirming specific complex formation between SA–anti-P4 and gA–P4.

**Fig. 4.**
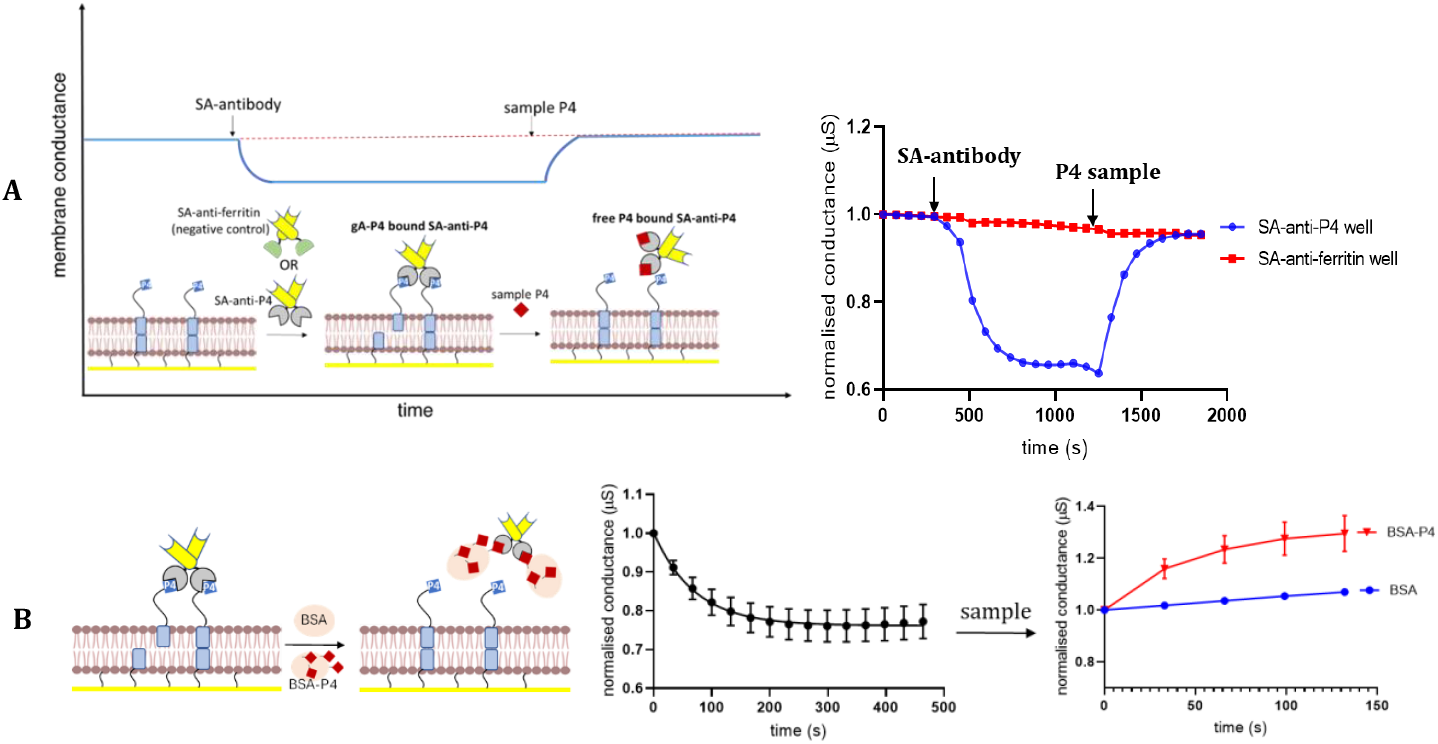
(A): Analyte detection using the SA-antibody proteins and tBLM containing gA-P4 molecules. Addition of SA-anti-P4 resulted in a decrease in the membrane conductance. A subsequent addition of P4 analyte sample resulted in an increase in the membrane conductance. No signals were detected for the control group (SA-anti-ferritin group). A rapid and specific strategy for detection of free analyte achieved in less than 15 minutes including SA-antibody and free analyte addition steps. The system allows for single step wash-free competitive analyte detection. (B): Specific detection of P4 conjugated to BSA using the SA-anti-P4 proteins and tBLM containing gA-P4 molecules. Addition of SA-anti-P4 resulted in a decrease in the membrane conductance. A subsequent addition of BSA-P4 analyte sample resulted in an increase in the membrane conductance. No signals were detected for the control analyte group (BSA) (n=3), (P = 0.0201).

As shown in Fig. 4, no washing steps were required, demonstrating the feasibility of a rapid (<15 min), wash-free ICS assay for both small (P4) and large (BSA-P4) analytes. For small molecules such as P4, a competitive switch mechanism enabled specific detection using a simple, label-free, and reagent-less strategy.

Wash-free biosensors are highly suited for point-of-care diagnostics due to their speed and operational simplicity. However, existing platforms face limitations: fluorescence-based methods are susceptible to background interference, require complex instrumentation, and involve lengthy assay times (Kaur et al., 2022), nanoparticle-based systems can aggregate and typically provide only qualitative results; electrochemical systems may lack selectivity and are environmentally sensitive (Huang et al., 2017). Protein-engineered biosensors, while promising, often involve complex and time-consuming production (Adamson et al., 2019). In comparison, ICS biosensing— particularly when coupled with recombinant binders—offers a clean, label-free alternative with minimal reagent requirements. The use of engineered binders allows for the design of competitive or non-competitive assay formats. ICS avoids optical interference due to its electrical readout, and benefits from the anti-fouling properties of tethered lipid membranes (Li et al., 2021; Mckeating et al., 2019), enabling analyte detection in unprocessed samples. Furthermore, ICS assay times are comparable to or faster than many existing wash-free systems. ICS supports both competitive detection of analytes with defined paratopes and non-competitive detection of multimeric or associated binders, such as SA-anti-P4 (Fig. 3C). This dual-mode detection, together with its wash-free, recombinant binder-enabled format, highlights ICS as a promising platform for future biosensing applications.

#### 3.4.2. Molecular Mechanism of Competitive Analyte Detection Using ICS and Recombinant Binding Proteins

In ICS, competitive detection is based on the displacement of recombinant analyte-binding proteins pre-associated with gA-modified lipid membranes. These binders recognise the target analyte or its conjugate, blocking channel dimer formation and reducing conductance. Upon introduction of free analyte, competition for the binder leads to its release, restoring conductance. This mechanism allows rapid, label-free detection of both small and large analytes, as demonstrated with P4 and BSA-P4 using SA-anti-P4. Molecular docking simulations (Supplementary Figure 2) further supported this system, showing that SA-anti-P4 binds more strongly to free P4 than to its conjugated form, which aligns with our experimental results.

#### 3.4.3. Quantitative Interaction Detection

Following the development of the competitive, wash-free biosensing format, the platform was evaluated for its ability to quantify analyte concentrations using the recombinant analyte binder SA-anti-P4 (principle of the quantitative signal analysis depicted in Supplementary Figure 3). Figures 5 to 7 illustrate the quantitative detection of analytes enabled by the recombinant analyte-binding protein.

**Fig. 5.**
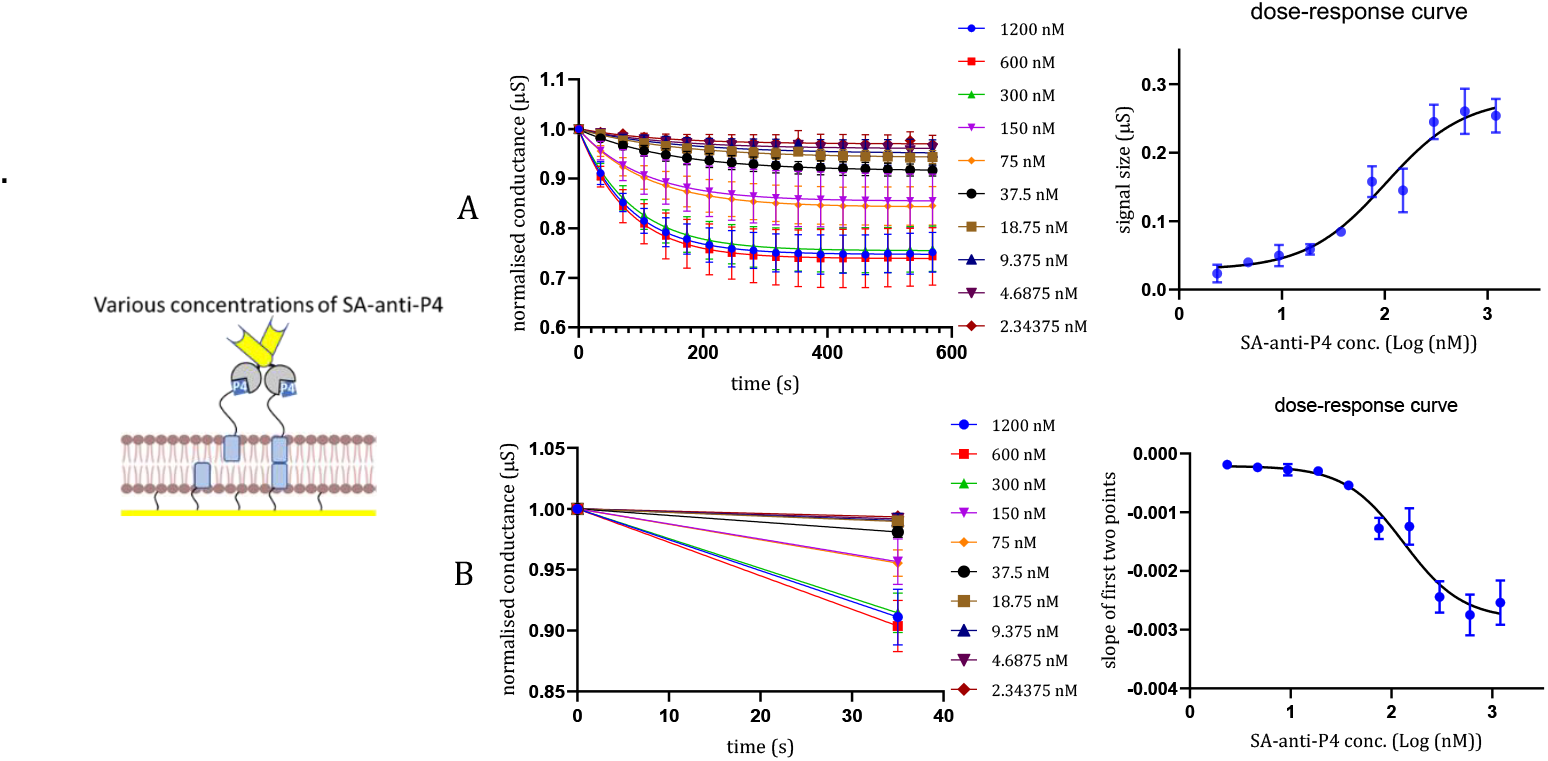
Quantitative detection of SA-anti-P4 (n=3). (A): the equilibrium or span based detection which took around 600 seconds versus (B): pre-equilibrium detection of SA-anti-P4 binding by calculation of slope from first two time points that took around 35 seconds. The analysis dose-response curve in both cases showed a similar linear response spanning 18.75 to 600 nM range.

**Fig. 6.**
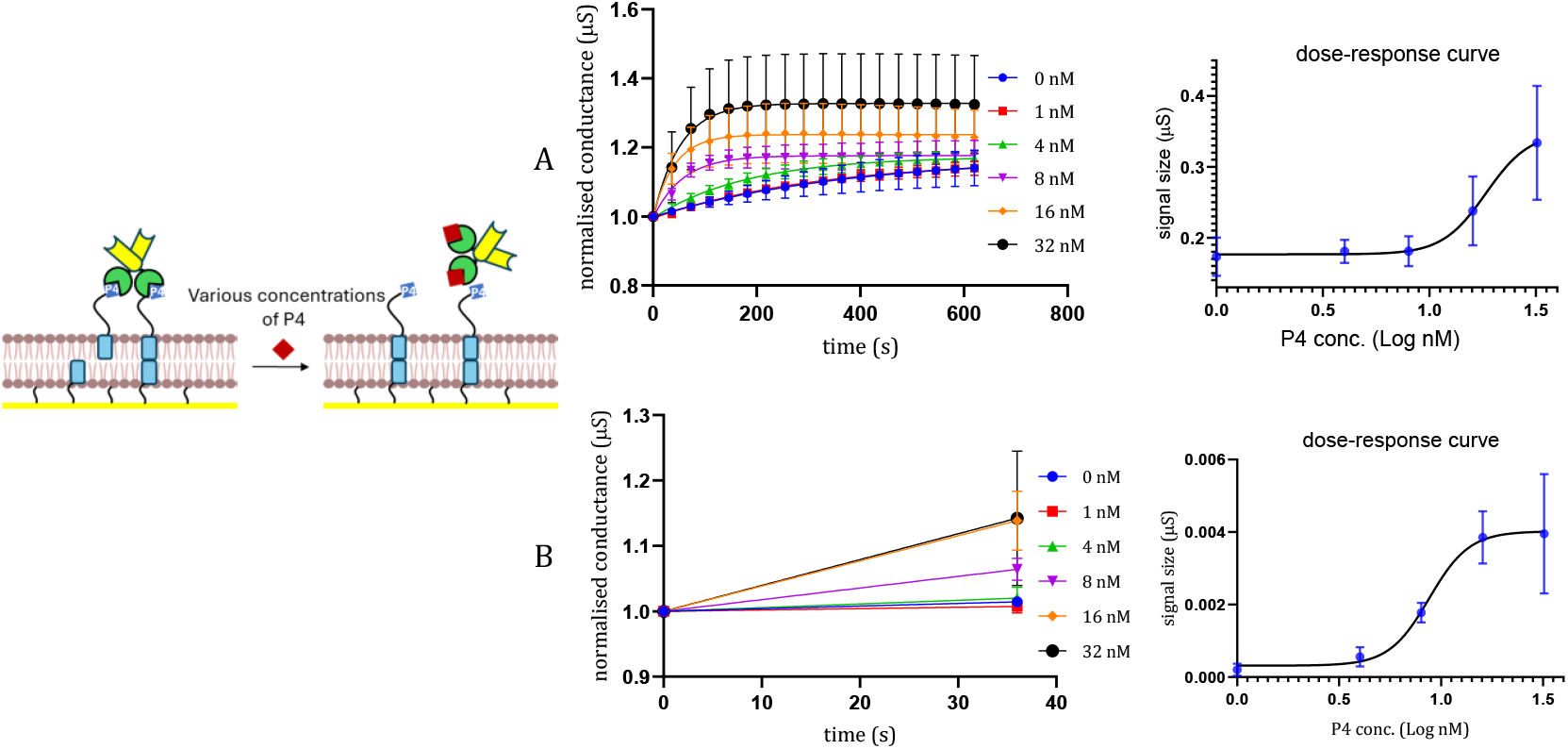
Quantitative detection of P4 (n=3). (A): the equilibrium or span based detection which took around 600 seconds with a linear range of 8-32 nM. (B): pre-equilibrium detection quantitation by calculation of slope from first two time points that took around 35 seconds with linear range of 4-16 nM.

**Fig. 7.**
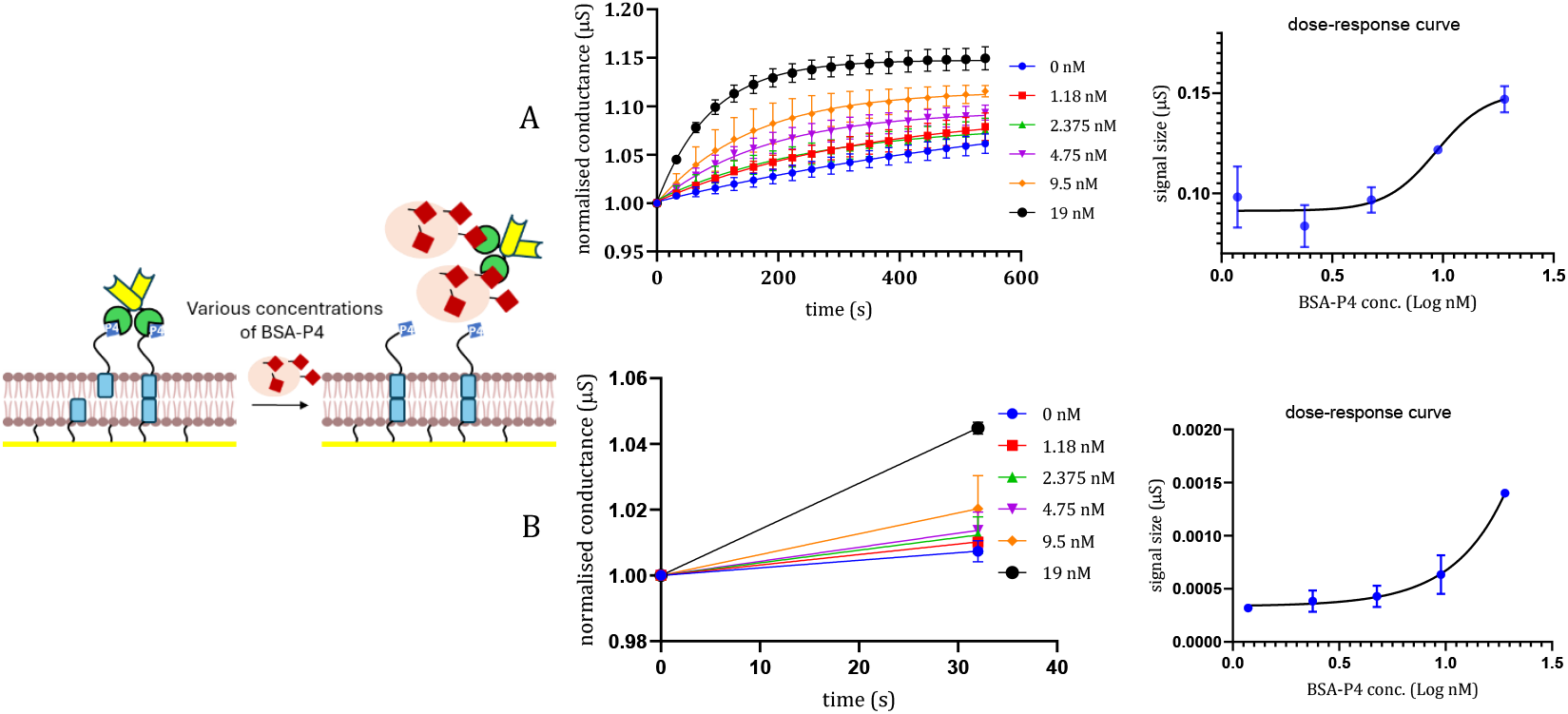
Quantitative detection of BSA-P4 (n=3). (A): the equilibrium or span based detection which took around 600 seconds with a linear range of 4-19 nM. (B): pre-equilibrium detection quantitation by calculation of slope from first two time points that took around 35 seconds with linear range of 4-19 nM.

The synergy of recombinant protein engineering and the Ion Channel Switch (ICS) biosensor enables real-time, label-free, and quantitative detection of analytes in both competitive and non-competitive assay formats. The use of recombinant components enhances assay performance while supporting a modular platform for rapid diagnostic development.

A genetically engineered single-chain variable fragment (scFv), SA-anti-P4, was designed to specifically recognise P4. In a non-competitive format, this recombinant antibody directly detected binding to a gA-P4-functionalised lipid membrane without secondary reagents or labelling. The ICS biosensor registered marked conductance shifts upon SA-anti-P4 binding, confirming its role as a primary detection molecule. The limit of detection (LOD) was determined to be 6 nM in pre-equilibrium and 12 nM at equilibrium, with limits of quantification (LOQ) of 31.6 nM and 36.45 nM, respectively, demonstrating the suitability of recombinant scFvs for sensitive, direct analyte detection using ICS systems.

The analyte-interacting element of the ICS, gramicidin A conjugated to P4 (gA-P4), functioned both as an ion channel and an epitope-presenting scaffold. This construct formed functional channels in lipid bilayers while exposing the P4 moiety for specific binding. Upon interaction with SA-anti-P4, the biosensor exhibited dose-dependent conductance changes with a linear dynamic range between 18.75 and 600 nM.

Real-time quantification of analyte binding was achieved by analysing conductance slopes between early time points, bypassing the need for equilibrium. This approach reduced assay times to under 60 seconds while maintaining or improving sensitivity. In competitive formats, detection of free P4 showed a pre-equilibrium LOD of 3.7 nM (vs. 5.14 nM at equilibrium), and detection of BSA-conjugated P4 (BSA-P4) reached 8.17 nM (vs. 13.64 nM).

The same recombinant reagents were applied across both assay formats. In the competitive setup, free P4 or BSA-P4 competed with SA-anti-P4 for gA-P4 binding on the membrane, resulting in signal attenuation. The interchangeable use of components without re-optimisation highlights the modularity of the system and its adaptability for a range of target types including small molecules, peptides, and proteins.

The biosensor’s lipid bilayer structure closely mimics biological membranes, allowing recombinant membrane-integrating proteins like gA-P4 to retain their native conformation and activity. This structural relevance improves assay accuracy, particularly for targets that are membrane-associated or conformation-sensitive. The integration of such constructs marks a significant advancement over traditional platforms.

Altogether, the combination of recombinant scFvs and engineered ion channel peptides within a biomimetic ICS platform establishes a new direction in biosensing—ultrafast, label-free, and modular within a physiologically relevant environment. This approach supports rapid assay development and is well suited for diverse diagnostic applications across biomedical and environmental fields.

#### 3.4.4. Using the protein in wash-free analyte detection: Harnessing bispecific interaction for continuous analyte detection

To explore continuous biosensing, the bispecific SA-anti-P4 protein, containing both streptavidin and anti-P4 scFv domains, was utilised to enable repeated analyte detection without the need for re-addition of protein. This was tested using tethered bilayer lipid membranes (tBLMs) incorporating both gA-P4 and gA-biotin channels, designed to retain the recombinant binder on the membrane surface throughout multiple detection cycles.

P4 was detected repeatedly in the same well using this setup. Control tBLMs with only gA-P4 and test membranes with a mixture of gA-P4 and gA-biotin were compared. As shown in Fig. 8, the mixed membrane allowed for multiple rounds of P4 detection, with the signal regenerating after each rinse. Rinsing removed residual P4 from the well, permitting SA-anti-P4 to rebind to gA-P4 channels and re-establish a responsive surface.

**Fig. 8:**
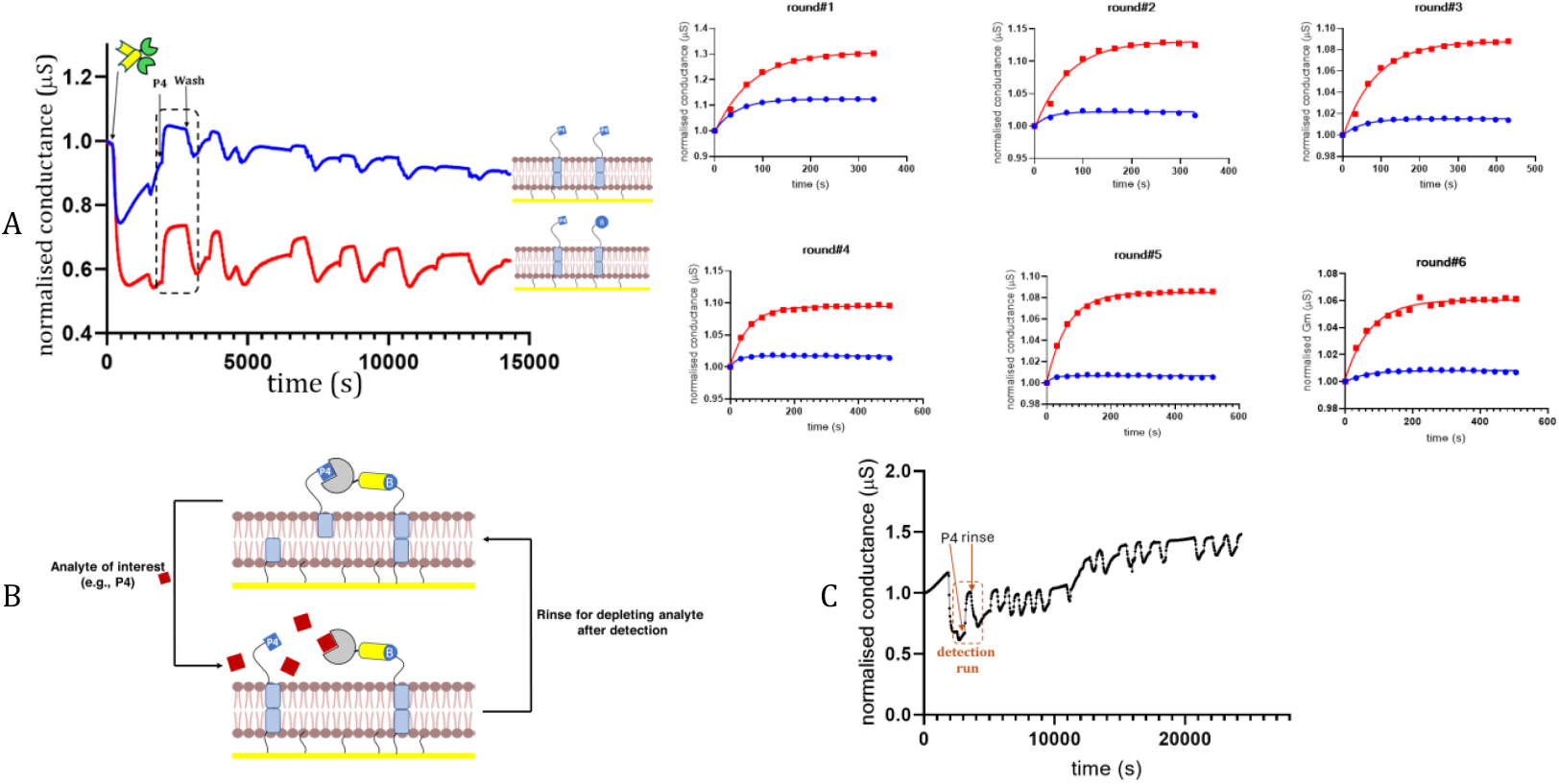
Repeated detection of P4 using mixed second layer. (A): The mixed second layer of gA-P4 and gA-biotin enabled continued detection of 10 nM P4 sample. After cycle 6, P4 response was almost undetectable when only gA-P4 was used in the second layer. (B): The suggested mechanism for the continuous detection of analytes using competitive P4 ICS biosensing. In a tBLM with gA-P4 and gA-biotin in the second lipid leaflet, SA-anti-P4 binds to biotin and P4. Addition of P4 results in competition between the free added P4 and P4 conjugated to gA in binding to P4 binding pocket of SA-anti-P4, and due to difference in binding affinities of the unconjugated and conjugated P4 molecules the added P4 replaces conjugated P4, resulting in release of the upper leaflet gAP4. Subsequently, gAP4 monomer is free to diffuse and form conducting dimer with the bottom leaflet gA monomer. However, binding between SA domain and the gA-conjugated biotin is not affected, and SA-anti-P4 remains anchored to the membrane via binding to gABiotin. For doing the next round of detection, rinsing step is included to deplete the added/detected P4 from the biosensing surface which results in re-binding of anti-P4 domain to gAP4, and a subsequent gating signal. Multi-hour-long continuous detection of P4 molecule. (C): A seven-hour long experiment showed capability of P4 ICS in continuous detection of P4 using SA-anti-P4 and tBLM with gA-P4 and gA-biotin.

A long-term assay using 10 nM P4 confirmed that the sensor remained functional after 16 detection cycles across seven hours, supporting the feasibility of this approach for extended, continuous detection applications (Fig. 8C).

SA-biotin coupling was employed to anchor the bispecific SA-anti-P4 protein onto the membrane surface via biotin-gA channels, enabling a P4 ICS configuration with real-time, continuous detection capabilities. This strategy successfully demonstrated that the SA-anti-P4 construct can support repeated analyte sensing cycles within the same well, establishing a foundational model for continuous biosensing using small molecule targets.

Future studies may explore the covalent coupling of receptor proteins to membrane elements. Covalently anchored receptors could further extend the duration and robustness of continuous monitoring. The engineered anti-P4 scFvs used in this study may serve as a prototype for such permanently anchored receptors, facilitating the development of durable and regenerable biosensor platforms.

The capability for continuous analyte detection offers significant advantages, such as reducing overall assay time and cost, while enabling repeated measurements from a single biosensor. This is particularly beneficial in long-term monitoring experiments, where biosensor replacement is impractical. In the current study, the mixed layer tBLM format enabled signal regeneration across more than 15 detection cycles in nanomolar P4 samples, with the signal remaining clearly distinguishable throughout. This confirms the platform’s applicability to multi-hour monitoring scenarios and suggests adaptability to longer durations based on application needs.

The developed real-time, continuous ICS system for detecting P4, a small steroid molecule, demonstrates superior performance compared to other biosensing platforms. For example, Arroyo-Curra s et al. (Arroyo-Curra s et al., 2017) and Ferguson, B. S. et al. (Ferguson et al., 2013) developed an aptamer-based electrochemical biosensor for real-time, in vivo drug monitoring in awake mice. While innovative, these systems lack nanomolar sensitivity and is not portable (Downs and Plaxco, 2022; Ferguson et al., 2013), limiting its broader applicability.

Another method based on particle diffusion dynamics has been reported for monitoring small molecules such as creatinine (Yan et al., 2020) and cortisol (Buskermolen et al., 2022). This system relies on microscopy to track particle motion, which changes upon analyte binding. Despite its sensitivity and continuous monitoring capabilities, this method requires 3–13 minutes for response and depends on microscope-based detection systems (Buskermolen et al., 2022; Yan et al., 2020), making it less portable and more time-consuming than the ICS approach. Furthermore, its detection limit for cortisol remains in the micromolar range (Buskermolen et al., 2022), indicating lower sensitivity compared to the P4 ICS system.

A third strategy uses antibody-switch constructs, where antibody–DNA conjugates with fluorophores and FRET pairs detect digoxigenin and cortisol in real-time (Thompson et al., 2023). In this method, analyte binding alters FRET signals depending on molecular displacement. While this assay reaches detection limits of 3.3 nM for cortisol and 5.8 nM for digoxigenin with ∼10-minute response times, it involves multiple reagent components and complex optimisations. Additionally, its fluorescence-based readout is vulnerable to photobleaching and autofluorescence interference (Kaur et al., 2022), which limits its long-term usability.

In contrast, the P4 ICS system developed here integrates a genetically engineered bispecific protein with a regenerable membrane interface, offering nanomolar sensitivity, real-time monitoring, portability, and minimal reagent preparation. These features position the ICS platform as a versatile and powerful solution for continuous biosensing of small molecules, with strong potential for deployment in a range of research, clinical, and environmental monitoring applications.

## 4. Conclusions and Future Perspectives

This study established the utility of Ion Channel Switch (ICS) technology for the functional characterisation of engineered bispecific recombinant proteins. The ICS platform enabled the real-time, label-free, and wash-free detection of both small and large analytes, and effectively distinguished monomeric from multimeric forms. The gating effect observed with SA–anti-P4 likely reflects molecular association via streptavidin domain. Although this was not directly confirmed by native-state analysis, the ICS response supports this interpretation and will be explored further in future work through structural biology techniques.

At least two functional binding sites per binder were required to generate a detectable signal, confirming the importance of binding valency in signal transduction.

The successful gating response generated by SA-anti-P4 and SA-anti-ferritin binders, when combined with their respective gA-modified channels (gA-P4 and gA-biotin), highlights the platform’s potential for detecting associative forms of proteins, including disease-relevant biomarkers and anti-steroid autoantibodies.

Repeatable, multi-hour detection of P4 using a mixed-layer ICS format demonstrated its applicability to continuous monitoring scenarios. The specificity of the system was preserved across binder types, gA modifications, and sample backgrounds, supporting its broader use in quantitative analyte detection.

ICS provides a sensitive and portable platform suitable for therapeutic binder screening, diagnostic assay development, and bioprocess quality assessment. The characterisation of recombinant binders based on parameters such as binding valency and avidity offers a direct and efficient alternative to conventional methods. Future studies will explore the integration of recombinant protein–ICS systems into complex matrices to further advance their application in diagnostics and drug development.

## CRediT authorship contribution statement

**Mohammad Pourhassan Moghaddam:** Methodology, Investigation, Formal analysis, Data curation, Writing – original draft and revise. **Krishanthi Jayasundera:** Methodology, Investigation, Validation. **Lele Jiang:** Methodology, Investigation, Validation, Conceptualisation. **Bruce A. Cornell:** Writing – review & editing, Validation, Supervision, Funding acquisition, Formal analysis, Conceptualization. **Stella M. Valenzuela:** Writing – review & editing, Validation, Supervision, Funding acquisition, Formal analysis, Conceptualization.

## Acknowledgements

This work was supported by the Australian Research Council (ARC) Industrial Transformation Research Hub for Integrated Devices for End-user Analysis at Low-levels (IDEAL) (IH150100028). The authors thank Surgical Diagnostics Pty Ltd for their scientific contributions. M.P.M. was supported by the Australian Government Research Training Program.

## Declaration

A preliminary version of part of this work was presented in abstract form (Pourhassan Moghaddam et al., Abstract #122, The 48th Lorne Conference on Protein Structure and Function 2023), focusing on monomer-multimer discrimination using ICS. The current study expands this by applying ICS to wash-free, real-time quantification and continuous biosensing using engineered recombinant binders.

## Data availability

Data are available upon request.

