## supplementary figures 1-3 for "Functional Characterisation of Recombinant Proteins Using Ion Channel Switch Technology: A Label-Free, Wash-Free Platform for Biotechnological Applications"

[$] **Current Affiliations:**

1. Graduate School of Biomedical Engineering, Faculty of Engineering, University of New South Wales, Sydney, NSW, Australia.
2. BioPoint Pty Ltd, Belrose, NSW, Australia.

[*] **Corresponding Author:**

Dr Mohammad Pourhassan Moghaddam

### Supplementary Figure 1

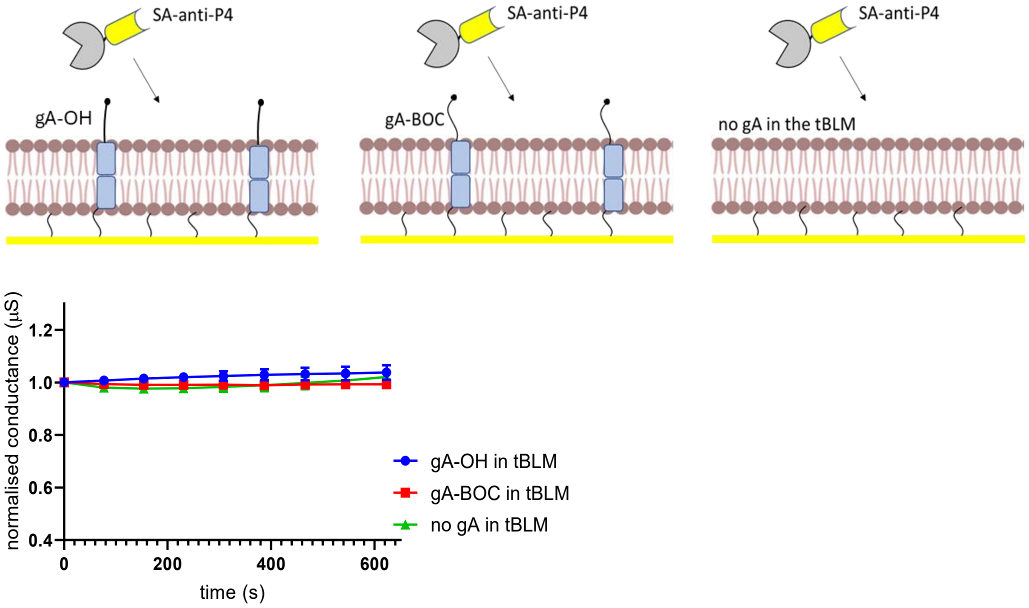


**Supplementary Figure 1.** Response of control tBLMs to SA-anti-P4 protein. To study non-specific bindings, three different control tBLM were used (n=3): tBLM containing non-modified gA molecules (gA-OH as an example of a gA molecule with chemically active end, and gA-BOC as an example of a gA molecule with chemically non-active end), and a tBLM without no gA molecule. The results showed no detectable decrease in the membrane conductance after addition of SA-anti-P4 protein, indicating no non-specific binding to the tBLM or irrelevant gA molecules.

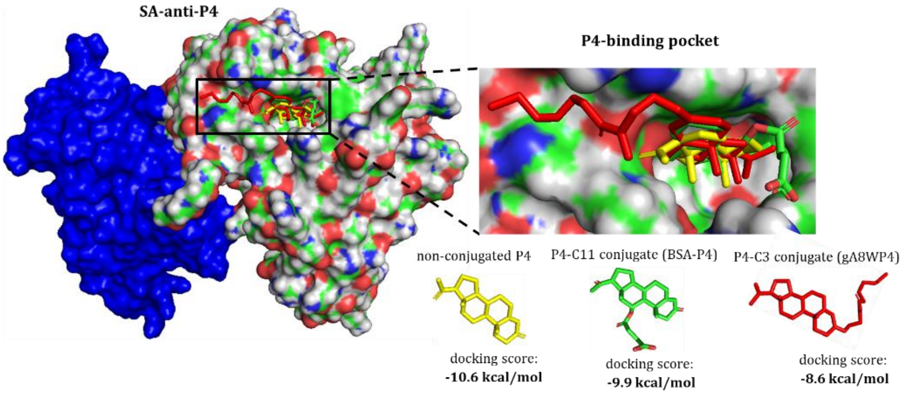


**Supplementary Figure 2.** Molecular docking of SA-anti-P4 protein with non-conjugated and conjugated P4 molecules. The non-conjugated P4 exhibited a lower docking score compared to the P4-C11 and P4-C3 conjugates, which are attached to BSA and gA8W, respectively. These results support the observed analyte competition in ICS assays.

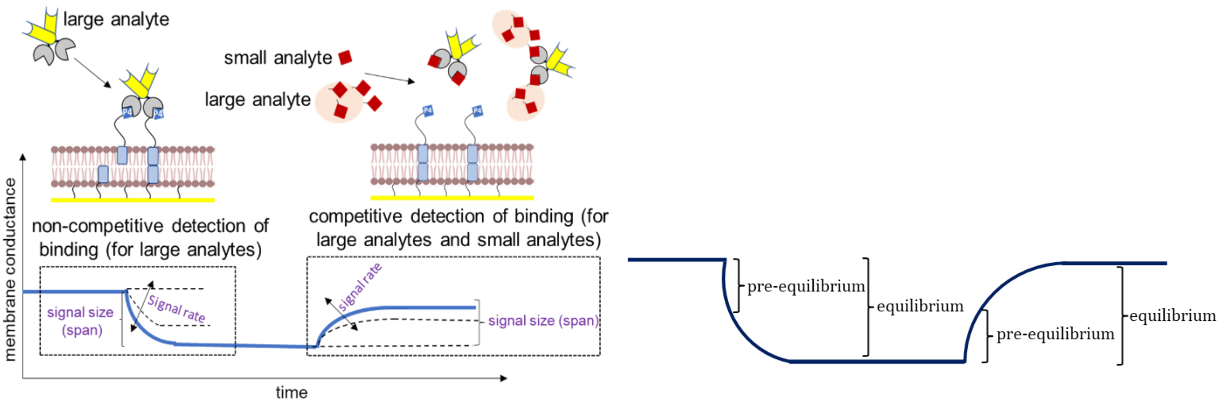


A B

**Supplementary Figure 3.** Quantitative detection of analytes by ICS based on non-competitive or competitive binding. (A): depiction of signal quantitation based on signal size and signal rate. Signal size or span is defined as the new level of membrane conductance after analyte binding. Signal rate is defined as speed of change in membrane conductance level after analyte binding, which can be the basis for kinetics analysis of binding events (B): equilibrium versus pre-equilibrium signal detection or quantitation. Equilibrium is the maximum level of signal size which stays steady after analyte addition, and preequilibrium is defined as signal change that is not reached at maxima point or pre-equilibrium point.
